# Biliverdin Reductase Catalytic Activity Is Essential for Malaria Resistance

**DOI:** 10.64898/2025.12.19.694840

**Authors:** Miguel Mesquita, Ana Figueiredo, Sonia Trikha Rastogi, Susana Ramos, Sara Pagnotta, Ana Rita Carlos, Silvia Cardoso, Ana Nóvoa, Moises Mallo, Bindu Paul, Elisa Jentho, Miguel P. Soares

## Abstract

Jaundice, a condition characterized by elevated levels of circulating bilirubin, is an adaptive response to malaria sustained through bilirubin production by biliverdin reductase A (BVRA). Beyond its enzymatic activity, BVRA acts as a protein kinase and as a transcription factor. To disentangle the contribution of BVRA catalytic activity over its non-canonical functions we generated *Blvra^G17A^* and *Blvra^E97A^* mice harboring G17A and E97A missense mutations in the BVRA NAD(P)H-binding domain and reductase motif, respectively. Both *Blvra^G17A^* and *Blvra^E97A^*mice presented a reduction in enzymatic activity and succumbed to malaria, otherwise non-lethal to wild-type (*Blvra^WT^*) mice. Quantification of circulating unconjugated bilirubin revealed a dose response effect whereby the mutant strains failed to reach a threshold of circulating bilirubin required to support its protective effect. These findings establish the antimalarial effect of the enzymatic activity of BVRA and define a concentration threshold of bilirubin required for malaria protection, informing therapeutic development and biomarker-guided malaria treatment strategies.

**Highlights:**

- Establishment of catalytic deficient BVRA mouse mutants;
- Anti-malarial effect of BVRA relies on its catalytic activity;
- A minimal bilirubin threshold for parasite control;
- A minimal bilirubin threshold for malaria resolution;

## Introduction

BVRA catalyzes the reduction of biliverdin IXα, an end-product of heme catabolism, into bilirubin IXα [1–3], a lipophilic radical-trapping antioxidant (RTA) and superoxide scavenger [4–6] to confer protection against oxidative stress [5–8]. However, BVRA also exerts “non-canonical” roles independent of its enzymatic activity [2; 9–14]. These have been attributed to a dual-specificity kinase (Ser/Thr and Tyr) and a DNA/chromatin-binding motif, at the N-terminal (aa 1-107) and C-terminal (aa 107-296) domains, respectively [2; 11; 15]. The dual kinase motif of BVRA phosphorylates the insulin receptor substrate-1 (IRS-1), thereby regulating insulin receptor signaling and downstream components of the MAPK/ERK pathway [16; 17]. Additionally, BVRA’s DNA/chromatin-binding activity modulates gene expression by activating transcription factor 2 (ATF-2) [10; 18]. More recently, BVRA was shown to exert cytoprotective effects independently of its catalytic activity, through regulation of the nuclear factor erythroid-derived factor-like 2 (NRF2) [19].

Circulating bilirubin is conjugated to glucuronic acid in the liver, via a reaction catalyzed by UDP glucuronosyltransferase family 1 member A1 (UGT1A1) [20], to prevent its accumulation over a cytotoxic threshold [21]. Only the conjugated bilirubin can be excreted through the bile and therefore when bilirubin conjugation falls below its production rate, unconjugated hyperbilirubinemia develops eventually leading to jaundice, a condition recognized clinically by yellowish discoloration of the skin and sclerae [22].

Jaundice is widely used clinically as a biomarker of liver dysfunction. While accurate, this led to the perception of jaundice representing a maladaptive response. The finding that unconjugated bilirubin was co-opted as a resistance mechanism against malaria calls for a reappraisal of jaundice [23].

The antimalarial effect of jaundice is attributed to the cytotoxic effect of unconjugated bilirubin over *Plasmodium spp.* parasites, the causative agents of malaria, suggesting that BVRA enzymatic activity is protective against malaria [19]. Here, we tested this hypothesis in mice carrying genetic disruption of BVRA catalytic activity, through the introduction of G17A or E97A missense mutations impairing the NAD(P)H-binding and reductase domain, respectively. We found that both mutations compromise malaria survival, suggesting that the antimalarial effect of BVRA relies strictly on its enzymatic activity. Comparison between mouse strains carrying G17A or E97A mutations and wildtype mice, revealed the minimum level of bilirubin required for its anti-malarial effect.

## Results

### CRISPR-Cas9-mediated generation of BVRA catalytic mutant mice

To determine whether BVRA’s protective effect against malaria relies strictly on its enzymatic activity, we generated two independent C57BL/6J mouse strains carrying a G17A mutation disrupting the NAD(P)H-binding motif (Fig. 1A, B) or a E97A mutation impairing the oxireductase biliverdin-binding motif of BVRA (Fig. 1F, G). By targeting these separate domains, we set to distinguish between cofactor-[NAD(P)H] and substrate (biliverdin IXα)-dependent mechanisms underlying BVRA’s anti-malarial activity.

**Figure 1.**
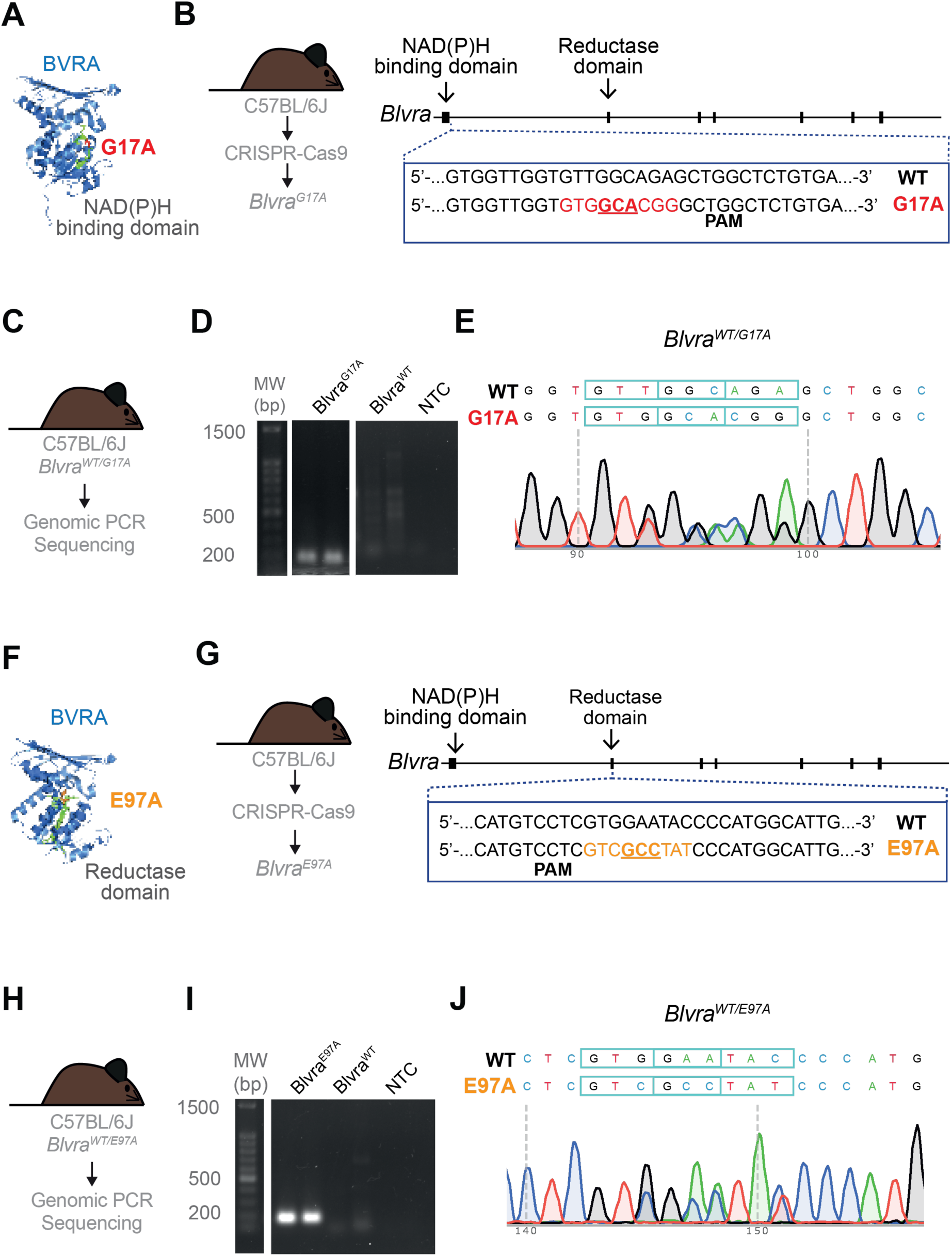
Generation And Validation Of *Blvra*^G17A^ and *Blvra*^E97A^ Mutant Mice. **(A)** Crystal structure of rat biliverdin reductase A (BVRA, PDB: 1LC3) showing the G17A mutation site within the NADPH-binding motif. The protein backbone is shown as a marine blue cartoon. The NADPH-binding motif (residues 13-19, green sticks) contains the conserved Rossmann fold (GXGXXG) essential for cofactor binding. The G17A mutation site is highlighted in red sticks, a critical position within this cofactor-binding motif. **(B)** Schematic representation of CRISPR-Cas9 targeting strategy showing the guide RNA (gRNA) target sequence and protospacer adjacent motif (PAM) for wild-type (WT) and mutant sequences to introduce the G17A mutation in the NAD(P)H-binding motif. **(C)** Schematic of genotyping PCR (gPCR) and Sanger sequencing analysis. **(D)** gPCR analysis showing gel electrophoresis of amplification products from *Blvra*^G17A^ mutant, *Blvra*^WT^ control samples and no template control as NTC, together with molecular weight (MW) markers indicated in base pairs (bp). **(E)** Sanger sequencing chromatogram confirming the precise insertion of the G17A mutation in *Blvra*^G17A^ mice compared to *Blvra*^WT^ controls. **(F)** BVRA crystal structure of rat biliverdin reductase A (BVRA, PDB: 1LC3) showing the E97A mutation site within the biliverdin-binding (reductase) domain (residues 90-105, green sticks). The protein backbone is shown as a marine blue cartoon. The E97A mutation is highlighted in orange sticks, indicating its position within the substrate-binding region. **(G)** Schematic representation of CRISPR-Cas9 targeting strategy showing the gRNA target sequence and PAM for WT and mutant sequences to introduce the E97A mutation in the reductase domain. **(H)** Schematic of genotyping gPCR and Sanger sequencing analysis. **(I)** gPCR analysis showing gel electrophoresis of amplification products from *Blvra*^E97A^ mutant, *Blvra*^WT^ control samples and no template control as NTC, together with molecular weight (MW) markers indicated in base pairs (bp). **(J)** Sanger sequencing chromatogram confirming the precise insertion of the E97A mutation in *Blvra*^E97A^ mice compared to *Blvra*^WT^ controls.

The *Blvra^G17A^* strain carries a GGCèGCA missense mutation replacing glycine (G) at position 17 by an alanine(A) (G17A) in the NADPH-binding domain of BVRA (Fig. 1A, B). This mutation is expected to disrupt the flexibility of the BVRA NADPH-binding loop [19], potentially affecting cofactor binding affinity and therefore catalytic efficiency (Fig. 1A, B). The GCA missense mutation was confirmed by genomic PCR (Fig. 1C, D) and validated by Sanger sequencing (Fig. 1C, E).

The *Blvra^E97A^* strain carries a GAAèGCC missense mutation replacing glutamate (E) at position 97 by an A (E97A) in the biliverdin-binding (reductase) domain of BVRA (Fig. 1F, G). This mutation is expected to disrupt BVRA oxireductase activity, impairing biliverdin IXα reduction into bilirubin IXα (Fig. 1F, G). The GCC missense mutation was confirmed by genomic PCR and Sanger sequencing (Fig. 1H-J).

### BVRA expression in *Blvra^E97A^* and *Blvra^G17A^* mouse strains

The relative level of *Blvra* mRNA expression in *Blvra^G17A^* mice was comparable to that of wild-type (*Blvra^WT^*) controls, as assessed by RT-qPCR in the spleen, kidney and liver (Fig. 2A). As expected, *Blvra^-/-^* mice showed neither *Blvra* mRNA nor protein expression (Fig. 2A,B). At the protein level, BVRA expression was lower in *Blvra^G17A^* mice, compared to the wild-type, as assessed by Western blot analysis (Fig. 2B). On the other hand, relative levels of *Blvra* mRNA expression were elevated in *Blvra^E97A^ vs. Blvra^WT^* controls, in the kidney and liver, but not the spleen, where there was a slight decrease as quantified by RT-qPCR (Fig. 2C). However, BVRA protein expression was only modestly reduced in the spleen of *Blvra^E97A^ vs. Blvra^WT^* controls, as assessed by Western blot analysis (Fig. 2D). These differences were not observed in kidney and liver extracts, which may reflect tissue-specific differences in protein levels (Fig. 2D).

**Figure 2.**
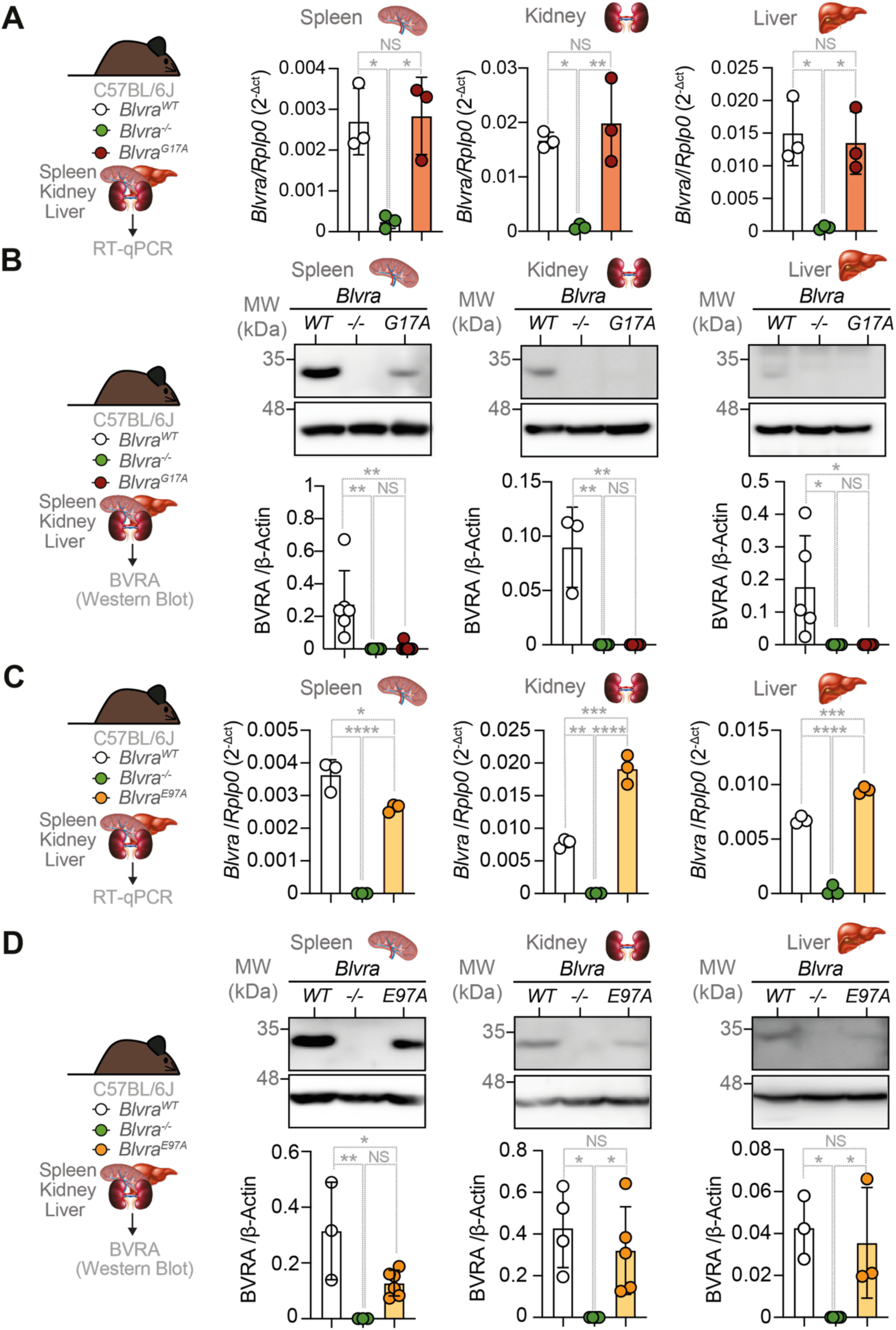
Expression Analysis of BVRA Missense Mutants G17A And E97A In C57BL/6J Mice. **(A)** qRT-PCR analysis of *Blvra* mRNA expression in spleen, kidney, and liver from *Blvra*^WT^, *Blvra^-/-^* and *Blvra^G17A^* mice. Data are expressed as *Blvra*/*Rplp0* (2^-ΔCt^) and presented as individual data points with bar graphs showing mean ± SD. n=3 mice *per* genotype. **(B)** Western blot analysis of BVRA protein expression in spleen, kidney, and liver. Representative blots show BVRA (33 kDa) and β-actin (42 kDa) loading control. Quantification of BVRA protein levels normalized to β-actin is shown below each blot as individual data points with bar graphs showing mean ± SD. At least n=3 per genotype. **(C)** qRT-PCR analysis of *Blvra* mRNA expression in spleen, kidney and liver tissues from *Blvra^WT^*, *Blvra^-/-^* and *Blvra^E97A^*mice. Data are expressed as *Blvra*/*Rplp0* (2^-ΔCt^) and presented as individual data points with bar graphs showing mean ± SD. n=3 per genotype. **(D)** Western blot analysis of BVRA protein expression in spleen, kidney, and liver tissue. Representative blots and quantification of BVRA protein levels normalized to β-actin below each blot, shown as individual data points with bar graphs showing mean ± SD. n=3 per genotype. Statistical comparisons were performed using one-way ANOVA with Tukey’s post hoc test. *p<0.05, **p<0.01, ***p<0.001, ****p<0.0001; NS, not significant.

### Bilirubin production is diminished in *Blvra^E97A^* and *Blvra^G177A^* mice

To assess the functional consequences of the E97A and G17A mutations on BVRA enzymatic activity, we performed *ex vivo* activity assays in liver, spleen, and kidney homogenates from *Blvra^E97A^*, and *Blvra^G17A^ vs. Blvra^WT^* and *Blvra^-/-^*mice. BVRA enzymatic activity was barely detectable in the spleen of *Blvra*^G17A^ mice, as compared to *Blvra^WT^* mice, where splenic BVRA enzymatic activity was estimated at ∼0.2 nM of bilirubin/μg protein/minute (Fig. 3A). Area under the curve (AUC) analysis confirmed a ∼95-97% reduction of enzymatic activity in the spleen of *Blvra*^G17A^ *vs. Blvra^WT^* controls, comparable to a 97-98% reduction in *Blvra*^-/-^ mice, with no significant difference between mutant genotypes (Fig. 3B). These results are consistent with loss of catalytic activity with the BVRA G17A mutant *in vitro* [19; 24].

**Figure 3.**
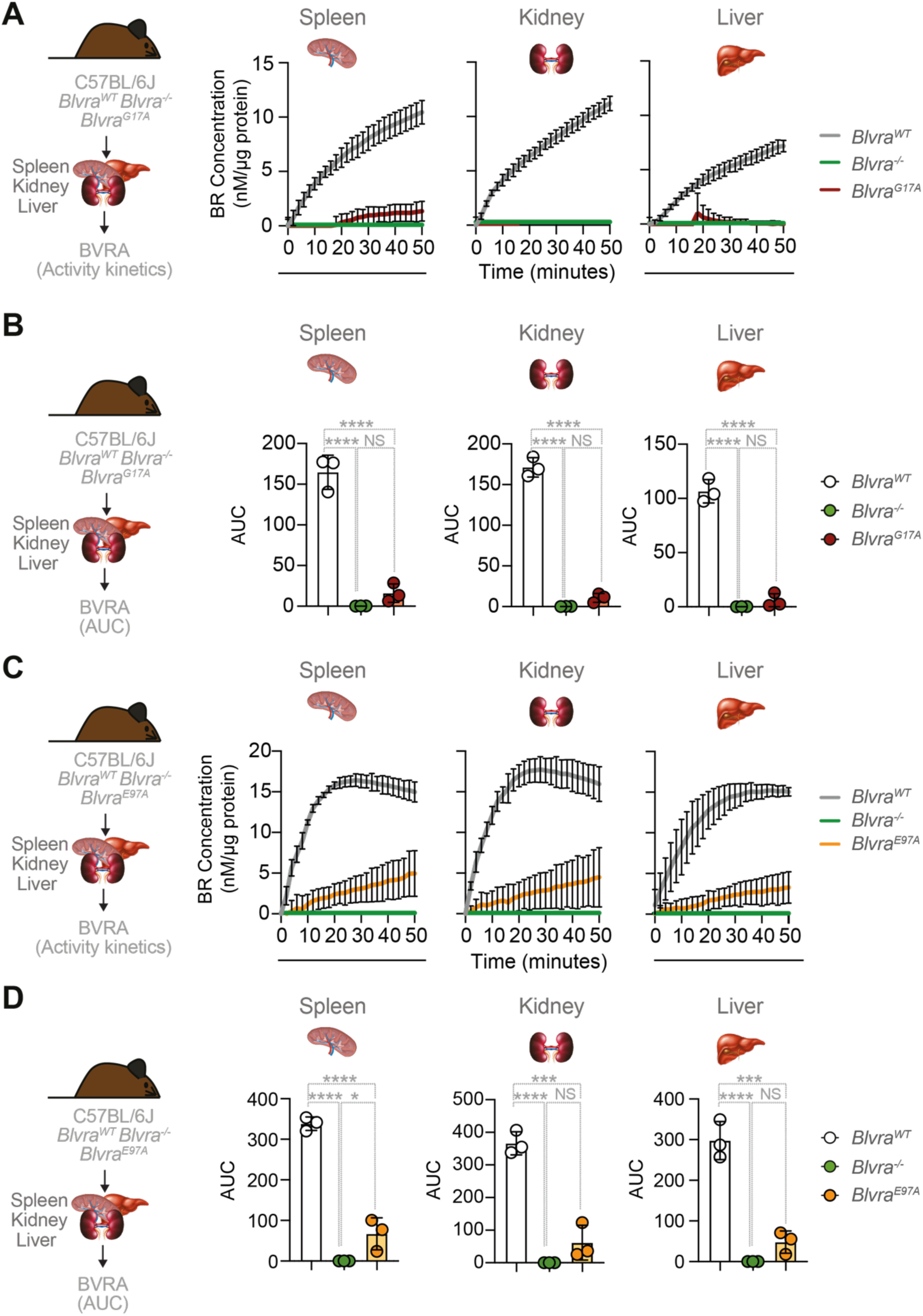
BVRA Enzymatic Activity Is Repressed In *Blvra*^G17A^ And *Blvra*^E97A^ Mutant Mice. (A-B) *Ex-vivo* BVRA enzymatic activity in tissue homogenates from *Blvra*^G17A^ mice. **(A)** Time course of bilirubin production measured as bilirubin concentration (nM/μg protein) over 50 minutes in spleen (*left*), kidney (*middle*), and liver (*right*) from *Blvra*^WT^*, Blvra*^-/-^, and *Blvra*^G17A^ mice. **(B)** Area under the curve (AUC) analysis spleen (*left*), kidney (*middle*), and liver (*right*) comparing enzymatic activity between genotypes. **(C-D)** *Ex-vivo* BVRA enzymatic activity in tissue homogenates from *Blvra*^E97A^ mice. **(C)** Time course of bilirubin production measured as BR concentration (nM/μg protein) over 50 minutes in spleen (*left*), kidney (*middle*), and liver (*right*) from *Blvra^WT^*, *Blvra*^-/-^, and *Blvra^G17A^* mice. **(D)** Area under the curve (AUC) analysis for spleen (*left*), kidney (*middle*), and liver (*right*) comparing enzymatic activity between genotypes. Data represent mean ± SD. Statistical comparisons were performed using one-way ANOVA with post hoc test. *p<0.05, ***p<0.001, ****p<0.0001; NS, not significant.

The kidneys from *Blvra^WT^* mice presented a BVRA enzymatic activity corresponding to ∼0.24 nM of bilirubin/μg protein/minute, whereas *Blvra^G17A^* mice and *Blvra*^-/-^ mice displayed no detectable enzymatic activity (Fig. 3A). AUC analysis estimates a reduction of ∼95-97% in *Blvra^G17A^* mice and greater than 97% in *Blvra*^-/-^ mice, compared to *Blvra^WT^*controls, with no significant difference between mutant genotypes (Fig. 3B).

BVRA enzymatic activity in liver was relatively lower, compared to spleen or kidneys from *Blvra^WT^* mice, corresponding to ∼0.14 nM/μg protein/minute (Fig. 3A). Liver from *Blvra*^G17A^ mice presented residual BVRA activity while those from *Blvra^-/-^* mice showed no measurable activity (Fig. 3A). AUC analysis estimates a ∼90-93% reduction of BVRA enzymatic activity in *Blvra*^G17A^ mice and greater than 95% in *Blvra*^-/-^ mice compared to *Blvra^WT^* controls, with no significant difference between mutant genotypes (Fig. 3B).

The catalytic activity of BVRA was substantially diminished, yet still detectable, in the spleen, kidneys and liver of *Blvra^E97A^* compared to *Blvra^WT^* controls (Fig. 3C). The spleen from *Blvra^E97A^* mice exhibited lower BVRA enzymatic activity, corresponding to ∼0.15 *vs.* 0.75 nM/μg protein/minute in *Blvra^WT^* mice in the first 20 minutes (Fig. 3C). AUC analysis estimated a ∼80% reduction in BVRA enzymatic activity, yet higher in *Blvra^E97A^* when compared to *Blvra*^-/-^ mice (Fig. 3D). Kidney BVRA activity in *Blvra^E97A^* mice was also decreased (∼0.2 nM bilirubin/μg protein/min *vs.* ∼0.85 nM/μg protein/min in *Blvra^WT^* in the first 20 min.) (Fig. 3C), a ∼90–95% reduction by AUC, similar to *Blvra^-/-^* mice (Fig. 3D). Liver from *Blvra^E97A^* mice also showed a marked reduction, compared to *Blvra^WT^* controls (0.15 nM bilirubin/μg protein/min *vs.* 0.6 nM bilirubin/μg protein/min, respectively, in the first 20 min) (Fig. 3C), ∼85–90% lower than *Blvra^WT^* mice, with no significant difference from the *Blvra^-/-^*genotype as estimated by AUC (Fig. 3D).

These findings suggest that the G17A mutation in the NADPH/NADH binding motif of BVRA results in nearly complete loss of function while the E97A mutation in the reductase motif of BVRA causes a severe reduction of BVRA enzymatic function.

### BVRA enzymatic activity is essential to survive malaria

To investigate the contribution of BVRA enzymatic activity to its antimalarial effect [23], *Blvra^G17A^*, *Blvra^E97A^* and *Blvra^-/-^* mice were infected with *Plasmodium chabaudi chabaudi* (*Pcc*) AS, a non-lethal strain to C57BL/6J genetic background, matched with *Blvra^WT^*control mice [25]. We monitored multiple parameters including a precise quantification of serum unconjugated bilirubin using a UnaG-based assay [26], parasitemia (% infected RBC), parasite burden (Nbr. of infected RBC/µL), disease scores, survival as well as body temperature, weight and glycemia changes, through a 15-day time course of infection.

At steady state, prior to infection, *Blvra^G17A^* mice had barely detectable levels of unconjugated bilirubin, similar to *Blvra^-/-^*mice (Fig. 4B-C), consistent with almost complete loss of BVRA expression (Fig. 2B) and enzymatic activity (Fig. 3A, B). In contrast, *Blvra^E97A^* mice presented levels of circulating unconjugated bilirubin at steady state similar to those of control *Blvra^WT^* mice, corresponding to ∼1-2 μM (Fig. 4A-C). This confirms that the BVRA E97A mutation retains protein expression (Fig. 2D) and residual enzymatic function (Fig. 3C,D), while the G17A mutation abolishes almost completely bilirubin production.

**Figure 4:**
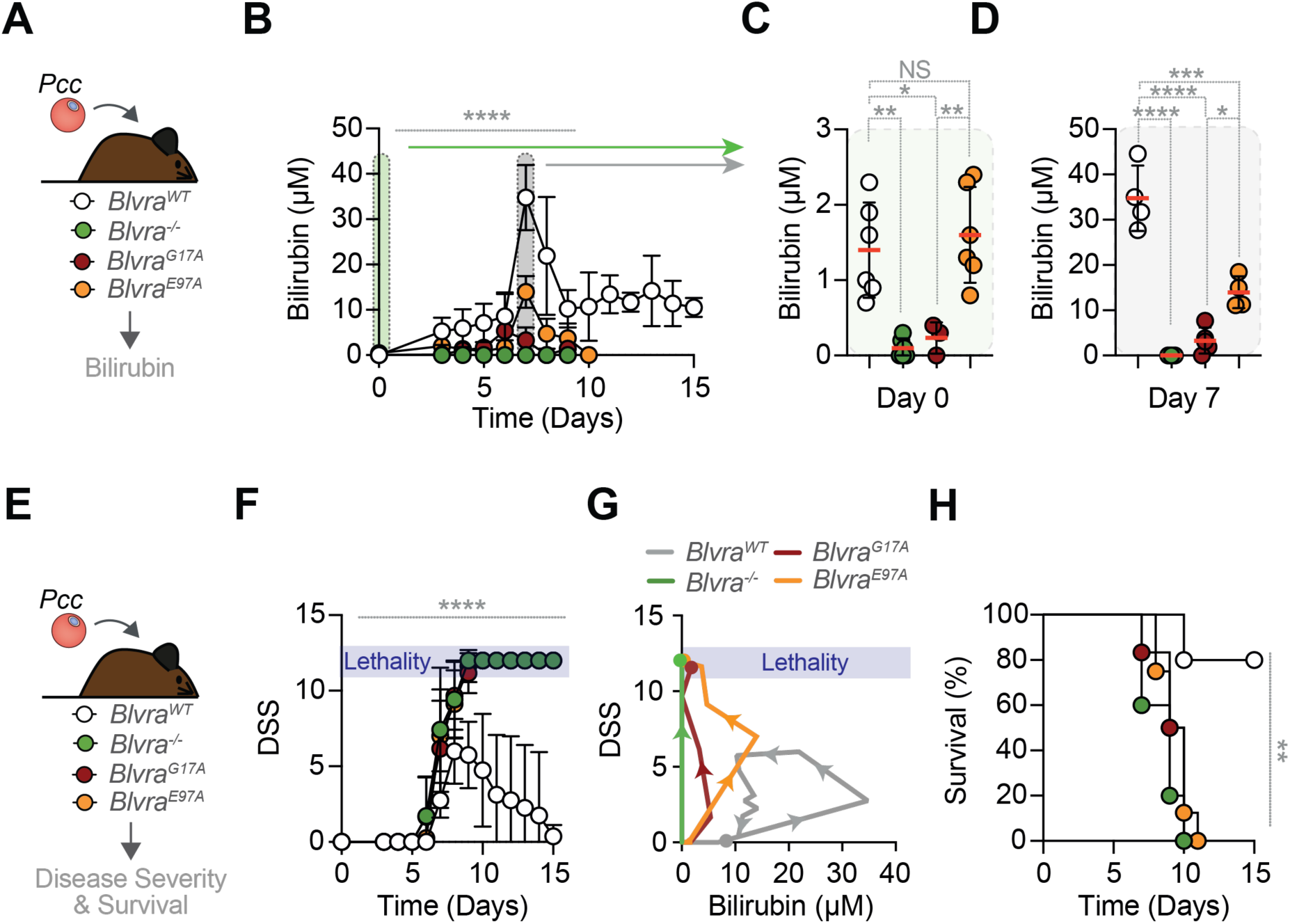
BVRA Catalytic Activity Determines Malaria Survival Through Dose-Dependent Bilirubin Production. **(A)** Experimental schematic showing *Plasmodium chabaudi chabaudi* (*Pcc*) infection in mice with different *Blvra* where bilirubin production was evaluated. **(B)** Time course of serum unconjugated bilirubin concentration following *Pcc* infection in *Blvra*^WT^, *Blvra*^-/-^, *Blvra^G17A^* and *Blvra^E97A^*mice (n = 5 to 8 per genotype) as measured by the UnaG-based assay. Data are shown as mean ± SD. **(C)** Serum unconjugated bilirubin levels before infection (green shaded region from panel B) in *Blvra*^WT^, *Blvra*^-/-^, *Blvra^G17A^* and *Blvra^E97A^* mice (n = 3 to 6 per genotype). **(D)** Serum unconjugated bilirubin levels at day 7 post-infection (gray shaded region from panel B) in *Blvra*^WT^, *Blvra*^-/-^, *Blvra^G17A^* and *Blvra^E97A^*mice (n = 5 to 8 per genotype). **(E)** Experimental schematic for disease severity and survival analysis. **(F)** Disease severity scores (DSS) over time following *Pcc* infection in *Blvra*^WT^, *Blvra*^-/-^, *Blvra^G17A^* and *Blvra^E97A^* mice (n = 5 to 8 per genotype). Data are shown as mean ± SD. **(G)** Disease trajectories showing the correlation between DSS and serum bilirubin levels for each genotype. Direction of disease trajectory is shown by arrows in each curve. **(H)** Kaplan-Meier survival curves of *Pcc*-infected *Blvra*^WT^, *Blvra*^-/-^, *Blvra^G17A^* and *Blvra^E97A^* mice (n = 5 to 8 per genotype). (A-H) Data pooled from n=3 independent experiments. Statistical comparisons were performed using two-way ANOVA (B, F); one-way ANOVA with Tukey’s multiple comparison test (C, D); log-rank (Mantel-Cox) test (H). *p < 0.05; **p < 0.01; ****p < 0.0001; NS, not significant.

*Blvra^WT^* mice developed jaundice in response to *Pcc* infection (Fig. 4A,B), consistent with described [23; 27]. The concentration of circulating unconjugated bilirubin in *Blvra^WT^* mice increased progressively in the first few days post-infection, reaching maximal ∼30-40 μM concentrations by day 7 (Fig. 4A,B,D). In contrast, *Blvra^G17A^*mice presented residual levels while *Blvra^-/-^* mice had no detectable levels of circulating unconjugated bilirubin throughout the course of infection (Fig. 4A,B,D). In contrast however, *Blvra*^E97A^ developed mild jaundice in response to *Pcc* infection, reaching a maximal level of ∼10-15 μM by day 7, lower than infected *Blvra^WT^* mice (Fig. 4A,B,D).

Malaria severity diverged markedly between *Blvra^WT^ vs. Blvra^G17A^*, *Blvra^E97A^* or *Blvra^-/-^*mice (Fig. 4E-G) with *Blvra^WT^* mice restraining disease severity scores (DSS) below 5 around days 7-9 and subsequently resolving DSS to 0 (Fig. 4E-G). In contrast, *Blvra^G17A^* and *Blvra^E97A^* mice developed maximal DSS scores of 10-13 by day 7-9 (Fig. 4E-G), similar to *Blvra^-/-^* mice (Fig. 4E-G).

The disease trajectories [28–30] established by the relationship of DSS and circulating unconjugated bilirubin, revealed a counter clock trajectory (Fig. 4G). *Blvra^WT^* reached a maximal unconjugated bilirubin of ∼35-40 µM concurrent with moderate ∼5-7 DSS scores, followed by coordinated decreases in both parameters’ indicative of disease recovery (Fig. 4G). In contrast, *Blvra^G17A^* mice displayed a near-vertical trajectory reaching maximal DSS of ∼12-13, concurrent with bilirubin accumulation below 5 µM, above *Blvra^-/-^* mice. *Blvra^E97^*mice showed an intermediate disease trajectory whereby bilirubin concentration reached a maximal level of ∼15-20 µM, failing to restrain an upward progression of DSS towards ∼10-12. These disease trajectories demonstrate that functional BVRA enzymatic activity is required to sustain levels of circulating bilirubin that mitigate malaria severity, similar to observed in *P. falciparum* malaria [23].

Approximately 80% of *Blvra^WT^* mice survived *Pcc* infection, consistent with the non-lethal nature of the *Pcc* AS strain in the C57BL/6J genetic background (Fig. 4H). In sharp contrast, *Blvra^G17A^* and *Blvra^E97A^* mice presented a progressive incidence of mortality starting at 7-8 days and reaching 100% by days 10-12 post-infection, similar to *Blvra^-/-^*mice (Fig. 4H) [23]. This demonstrates that BVRA enzymatic activity exerts a major impact on *Plasmodium* virulence shifting malaria outcome from non-lethal into fatal.

### BVRA enzymatic activity exerts anti-plasmodial effects

The kinetics of parasitemia revealed distinct temporal patterns between genotypes (Fig. 5A,B). Initially, all genotypes exhibited similar dynamics through day 6 post-infection, reaching the peak parasitemia of ∼50-60% (Fig. 5A,B). While all *Blvra^WT^* mice cleared parasitemia by day 10, this was not the case for *Blvra^G17A^* and *Blvra^E97A^* mice, which presented a parasitemia of ∼30-50% at day 8 post-infection, similar to *Blvra*^-/-^ mice (Fig. 5A,B), compared to ∼15-20% in *Blvra^WT^* mice (Fig. 5C). This is consistent with the BVRA catalytic activity containing parasitemia.

**Figure 5:**
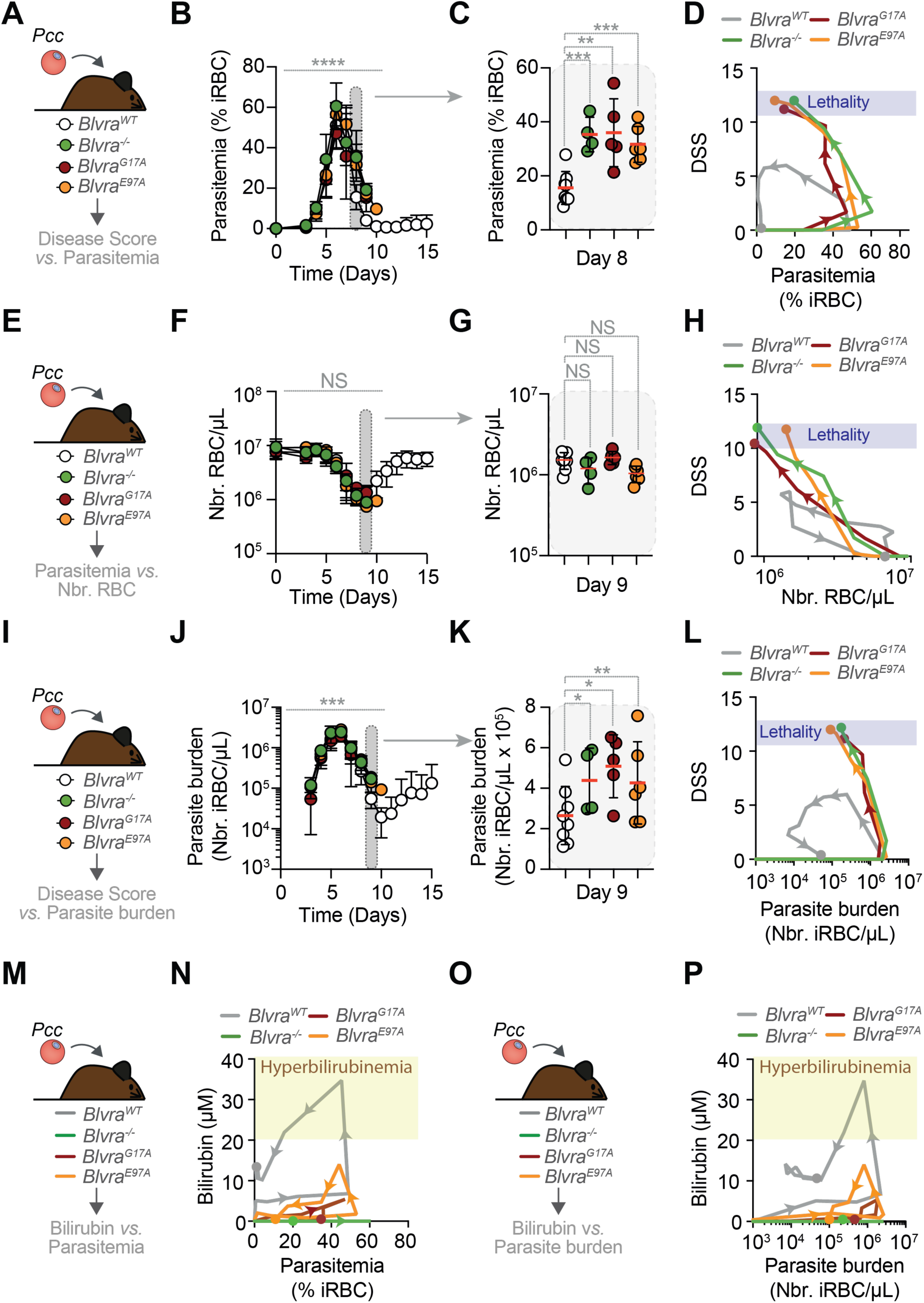
Biliverdin Reductase A Catalytic Activity is essential to limit life-threatening parasitemia. **(A)** Experimental schematic for disease severity and parasitemia. **(B)** Time course parasitemia (% infected RBC; iRBC) following *Pcc* infection in *Blvra^WT^*, *Blvra*^-/-^, *Blvra^G17A^*and *Blvra^E97A^* mice (n = 5 to 8 per genotype). Data are shown as mean ± SD. **(C)** Highlighted iRBC levels at day 8 (gray shaded region from panel B). Each circle represents an individual mouse. **(D)** Correlation between DSS and parasitemia for each genotype, showing inverse relationship in *Blvra*^WT^ mice. Direction of disease trajectory is shown by arrows in each curve. **(E)** Experimental schematic for total RBC analysis. **(F)** Time course of total RBC count following *Pcc* infection in *Blvra^WT^*, *Blvra*^-/-^, *Blvra^G17A^* and *Blvra^E97A^* mice (n = 5 to 8 per genotype). Data are shown as mean ± SD. **(G)** Highlighted total RBC count at day 9 (gray shaded region from panel F). Each circle represents an individual mouse. **(H)** Correlation between DSS and total RBC count for each genotype, showing inverse relationship in *Blvra^WT^*mice. Direction of disease trajectory is shown by arrows in each curve. **(I)** Experimental schematic for parasite burden analysis. **(J)** Time course of parasite burden (total infected RBC per μL blood; iRBC) following *Pcc* infection in *Blvra^WT^*, *Blvra*^-/-^, *Blvra^G17A^* and *Blvra^E97A^* mice (n = 5 to 8 per genotype). Data are shown as mean ± SD. **(K)** Highlighted parasite burdenlevels at day 9 (gray shaded region from panel J). Each circle represents an individual mouse. **(L)** Correlation between DSS and parasite burden for each genotype, showing inverse relationship most prominently in *Blvra^WT^*mice. **(M)** Experimental schematic for parasitemia and serum bilirubin. **(N)** Disease trajectories showing the correlation between parasitemia and serum unconjugated bilirubin levels for each genotype. Direction of disease trajectory is shown by arrows in each curve. **(O)** Experimental schematic for parasite burden and serum unconjugated bilirubin. **(P)** Disease trajectories showing the correlation between parasite burden and serum unconjugated bilirubin levels for each genotype. Direction of disease trajectory is shown by arrows in each curve. (A-P) Data pooled from n=3 independent experiments. Statistical comparisons were performed using two-way ANOVA (B, F, J); two-way ANOVA with Linear Mixed Effects Model (C, K) as primary analysis and treating independent experiments as linked experiments; one-way ANOVA with Tukey’s multiple comparison test (G) *p < 0.05; **p < 0.01; ***p < 0.001; ****p < 0.0001; NS, not significant.

The disease trajectories [28–30] established by the relationship of DSS *vs.* parasitemia showed again fundamental differences among genotypes (Fig. 5D). All genotypes exhibited counterclockwise loop trajectories, characterized initially by upward DSS inflection reaching ∼0-2 at peak parasitemia of ∼40-60% (Fig. 5D). While *Blvra^WT^* mice reached a maximal DSS of ∼5-6 transitioning towards full recovery (Fig. 5D), *Blvra^G17A^* and *Blvra^E97A^* mice presented an irreversible upward DSS progression to 13 (*i.e.,* death), similar to *Blvra*^-/-^ mice (Fig. 5D). This is consistent with the BVRA catalytic activity restraining malaria severity.

The development of anemia was comparable among *Blvra^WT^ vs. Blvra* mutant mice as monitored by the number of circulating RBC throughout the course of the infection (Fig. 5E,F). Total RBC counts at the peak of infection were similar across all mice (Fig. 5G), demonstrating that severe anemia does not account for the vulnerability of *Blvra* mutant mice during *Pcc* infection.

Analysis of disease trajectories established by the relationship of DSS *vs.* total RBC count demonstrate counterclockwise loop trajectories, characterized by an initial increase in DSS and decrease in total RBC count (Fig. 5H). *Blvra^WT^* mice developed maximal DSS of ∼5-6 at ∼2×10^6^ RBC/µL, transitioning to recovery from anemia, whereas *Blvra* mutant mice underwent irreversible DSS progression under the same levels of anemia (Fig. 5G).

Parasite burden followed kinetics similar to parasitemia (Fig. 5I,J). *Blvra^WT^* mice decreased parasite burden to approximately 2-3×10^6^ iRBC/µL at day 9 post-infection, while *Blvra^G17A^*and *Blvra^E97A^* mice presented 1.94- and 1.63-fold higher parasite burdens corresponding to ∼2-6×10^5^ iRBC/µL, respectively, similar to *Blvra*^-/-^ mice (Fig. 5K). This is consistent with the BVRA catalytic activity containing parasite burden at the peak of infection.

Analysis of disease trajectories [31] resulting from the relationship of DSS *vs.* parasite burden mirrored those of DSS *vs.* parasitemia (Fig. 5L). *Blvra*^WT^ mice maintained low DSS (<5), while *Blvra^G17A^* and *Blvra^E97A^* mice presented an irreversible upward DSS progression at parasite burdens exceeding 10^6^ iRBC/µL reaching DSS of 13 (*i.e.,* death). This is consistent with the BVRA catalytic activity exerting an antimalarial effect that contains malaria severity.

Disease trajectories representing the relationship of circulating bilirubin *vs.* parasitemia or parasite burden confirmed fundamental differences in *Blvra^WT^ vs. Blvra* mutant mice (Fig. 5M-P)). *Blvra^WT^*mice presented stable bilirubin concentrations at parasitemia below ∼50% (Fig. 5M,N) and parasite burdenof ∼2-3×10^6^ iRBC/µL (Fig. 5O,P). Above this threshold however, bilirubin concentration increased to reach a peak concentration of ∼30-35 µM, followed by a sharp decrease, associated with parasite clearance (Fig. 5M-P). In contrast, *Blvra^G17A^* and *Blvra^E97A^* mice failed to increase bilirubin and correspondingly, parasitemia and parasite burden were sustained, similar to *Blvra*^-/-^ mice(Fig. 5M-P), suggesting that the accumulation of bilirubin above ∼30 µM is necessary for effective parasite clearance.

### BVRA enzymatic activity regulates the host metabolic response to malaria

Plotting different host physiologic parameters against parasitemia and parasite burden also revealed fundamental differences in the disease trajectories of *Blvra^WT^ vs. Blvra* mutant mice (Fig. 6). Body temperature trajectories revealed profound thermoregulatory differences in *Blvra^WT^ vs. Blvra* mutant mice throughout infection (Fig. 6A-D). All genotypes displayed clockwise loop trajectories with stable body temperature (∼37°C) until a parasitemia/parasite burdenthreshold presenting a downward inflection towards hypothermia thereafter (Fig. 6A-D). While *Blvra^WT^* mice restrained hypothermia at sub-lethal levels (∼33-35°C) while reducing parasitemia and parasite burden(Fig.6A,B), this was not the case for *Blvra* mutants, which developed irreversible downward temperature trajectories culminating in life-threatening hypothermia (∼25-27°C)(Fig. 6A-D). The mutant strains showed overlapping disease trajectories, indicating equivalent thermoregulatory failure, consistent with BVRA catalytic activity conferring protection against life-threatening hypothermia during malaria (Fig. 6C,D).

**Figure 6:**
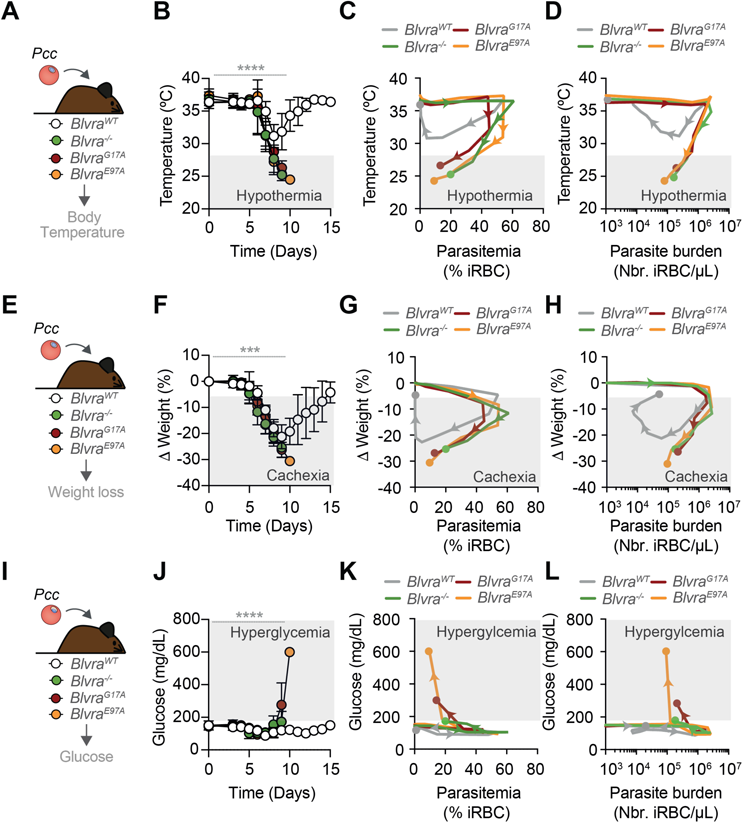
Biliverdin Reductase A Catalytic Activity Is Essential to Limit Severe Disease Outcome. **(A)** Experimental schematic for body temperature analysis during infection. **(B)** Body temperature over time following *Pcc* infection in *Blvra^WT^*, *Blvra*^-/-^, *Blvra^G17A^* and *Blvra^E97A^* mice (n = 5 to 8 per genotype). Data are shown as mean ± SD. **(C)** Correlation between body temperature and parasitemia for each genotype. Direction of disease trajectory is shown by arrows in each curve. **(D)** Correlation between body temperature and parasite burden levels for each genotype. Direction of disease trajectory is shown by arrows in each curve. **(E)** Experimental schematic for weight loss analysis during infection. **(F)** Weight loss over time following *Pcc* infection in *Blvra^WT^*, *Blvra*^-/-^, *Blvra^G17A^* and *Blvra^E97A^* mice (n = 5 to 8 per genotype). Data are shown as mean ± SD. **(G)** Correlation between weight loss and parasitemia for each genotype. Direction of disease trajectory is shown by arrows in each curve. **(H)** Correlation between weight loss and parasite burdenlevels for each genotype. Direction of disease trajectory is shown by arrows in each curve. **(I)** Experimental schematic for glucose analysis during infection. **(J)** Glucose levels over time following *Pcc* infection in *Blvra^WT^*, *Blvra*^-/-^, *Blvra^G17A^* and *Blvra^E97A^* mice (n = 5 to 8 *per* genotype). Data are shown as mean ± SD. **(K)** Correlation between glucose levels and parasitemia for each genotype. Direction of disease trajectory is shown by arrows in each curve. **(L)** Correlation between glucose levels and parasite burdenlevels for each genotype. Direction of disease trajectory is shown by arrows in each curve. (A-L) Data pooled from n=3 independent experiments. Statistical comparisons were performed using two-way ANOVA (E, I, M. ***p < 0.001; ****p < 0.0001.

Body weight trajectories throughout infection demonstrated metabolic differences between *Blvra^WT^ vs. Blvra* mutant mice analogous to those observed for thermoregulation (Fig. 6E-H). *Blvra^WT^* mice restrained weight loss (∼10-15%), whereas *Blvra^G17A^*, *Blvra^E97A^*and *Blvra^-/-^* mice underwent irreversible weight loss trajectories progressing towards severe cachexia (∼20-30%) (Fig. 6E-H). The overlapping trajectories of *Blvra* mutants are consistent with BVRA catalytic activity limiting metabolic dysfunction during malaria (Fig. 6G-H).

Glycemia trajectories confirmed major metabolic differences between *Blvra^WT^* mice *vs. Blvra* mutants (Fig. 6I-L). *Blvra^WT^*mice maintained stable glycemia (∼100-150 mg/dL) throughout infection, whereas *Blvra* mutants presented upwards glucose trajectories progressing severe hyperglycemia (∼400-600 mg/dL) at minimal parasitemia and parasite burden(Fig. 6I-L). Notably, *Blvra^E97A^* mice exhibited the most pronounced hyperglycemic response, consistent with BVRA catalytic activity controlling glycemia during malaria.

## Discussion

Through the generation and characterization of mouse strains carrying specific *Blvra* loss of function mutations we demonstrate that the antimalarial effect of BVRA relies essentially on its catalytic activity. This assumes that silencing the catalytic activity of BVRA does not interfere with its non-canonical functions [23], such as the regulation of NRF2 [19], a transcription factor that exerts a major impact on malaria severity [29; 32; 33].

The observation that the BVRA G17A mutation compromises BVRA protein expression despite physiologic mRNA expression levels in *Blvra^G17A^*mice (Fig. 2A,B), suggests that the glycine at position 17 within the NAD(P)H-binding domain confers conformational flexibility critical for cofactor binding and maintenance of proper protein folding. While previous protein circular dichroism (CD) analysis suggests that there are no changes in secondary structure of the BVRA G17A mutant [19], disruption of the NADPH binding motif in G17A may affect overall protein folding and stability *in vivo*, not fully captured by CD analysis of the purified protein [19]. It is possible therefore that the substitution of a glycine at position 17 by an alanine introduces steric constraints that destabilize folding of the NAD(P)H-binding pocket and could target BVRA for proteasomal degradation, presumably explaining the near-complete absence of BVRA protein despite normal transcript levels (Fig. 2A,B). This is consistent with the G17A mutation abolishing both enzymatic activity and protein stability, resulting in a phenotype functionally equivalent to *Blvra*^-/-^ mice (Fig. 3A,B, Fig. 4-6). In contrast, the BVRA E97A mutation severely impairs catalytic activity while still allowing for protein expression (Fig. 2C,D), suggesting that the NAD(P)H-binding domain supports cofactor recognition and structural integrity, whereas the reductase domain primarily governs catalytic function.

The observation that *Blvra^E97A^* mice maintain physiological bilirubin levels at steady state (Fig. 4A,B) is critical for mechanistic interpretation of the antimalarial effects of jaundice [23]. It demonstrates that E97A mutation is a catalytic hypomorph that allows circulating bilirubin to accumulate after birth, suggesting that the antimalarial effects of jaundice [23] are not related to a putative pre-sensitization mechanism due to baseline bilirubin deficiency. A possible metabolic adaptation to chronic bilirubin absence is not present in *Blvra^E97A^* mice and therefore the lethal outcome of malaria in *Blvra^E97A^* mice (Fig. 4H) should be attributed specifically to insufficient bilirubin production in response to *Plasmodium* infection.

The dose-response relationship across the *Blvra* mutant lines enables interpolation of the minimum effective concentration of unconjugated bilirubin providing antimalarial protection (Fig. 4A-D). *Pcc*-infected *Blvra^WT^* mice reached a maximal concentration of circulating unconjugated bilirubin in the ∼30-40 μM range, while *Blvra*^E97A^ mice accumulate no more than ∼10-15 μM, approximately 25-38% of *Blvra^WT^* mice (Fig. 4A-D). In contrast *Blvra^G17A^* mice maintain residual levels of bilirubin throughout infection (Fig. 4A-D). As *Blvra^E97A^* mice succumb to *Pcc* infection with kinetics nearly identical to *Blvra^G17A^* and *Blvra^-/-^*mice, this indicates that the protective threshold for unconjugated bilirubin lies between 20-30 μM. The convergence of a 20-40 μM unconjugated bilirubin concentration protective range in mice and 8-50 μM in asymptomatic humans [23], supports the idea of evolutionary conservation across host and *Plasmodium* species.

This dose-dependency is further supported by parasite clearance kinetics (Fig. 5). By day 8 post-infection, when *Blvra^WT^* mice begin resolving parasitemia, *Blvra*-mutant mice maintain significantly elevated parasitemia and parasite burdens (Fig. 5). The relative inability of *Blvra^E97A^* mice, with intermediate bilirubin levels, to achieve parasite control indicates a threshold phenomenon for the antiplasmodial effect of bilirubin, rather than linear dose-response. It suggests that below a threshold concentration, bilirubin cannot effectively achieve parasite clearance.

Mechanistically, bilirubin inhibits *P. falciparum* growth through a coordinated targeting of mitochondrial function [23; 34], inhibition of hemozoin crystallization and disruption of the parasite food vacuole [23]. The steep concentration-response relationship observed in the present study suggests that this antiplasmodial activity of bilirubin requires simultaneous disruption of multiple parasite processes, including: i) *de novo* pyrimidine synthesis catalyzed by mitochondrial dihydroorotate dehydrogenase (DHODH), glycolysis through inhibition of enolase, glucose-6-phosphate isomerase, phosphoglycerate kinase, phosphoglycerate mutase, ATP synthesis through inhibition of ATP synthase subunits, cytochrome c oxidase and hemoglobin digestion genes through inhibition of M18 aspartyl aminopeptidase, falcilysin, M17 leucyl aminopeptidase [23]. The threshold phenomenon likely reflects the minimum bilirubin concentration necessary to achieve antiplasmodial activity across these multiple targets simultaneously. At concentrations below 15 μM, individual mechanisms may be partially targeted, generating sub-lytic stress but the combined effect remains insufficient to arrest parasite development. At concentrations exceeding 20-30 μM, the cumulative impact on several metabolic pathways surpasses the parasite’s compensatory capacity.

The disease trajectory of *Blvra^WT^* mice *vs. Blvra* mutants provide a mechanistic framework for understanding how BVRA catalytic activity determines malaria outcomes. *Blvra^WT^* mice demonstrated both resistance (parasite clearance) and possibly tissue damage control [35; 36] mechanisms reducing disease severity (Fig. 4E-G), including hypothermia (Fig. 6A-D) and weight loss (Fig. 6E-H) as bilirubin exceeded 30 μM. In contrast, *Blvra^G17A^*, *Blvra^E97A^* and *Blvra^-/-^*mice sustained higher parasitemia with irreversible disease severity trajectories progressing toward life-threatening outcomes, revealing failed resistance. The superimposable trajectories of *Blvra* mutants across thermoregulation (Fig. 6A-D) and body weight (Fig. 6E-H) suggest that 20-30 μM bilirubin threshold operates through resistance rather than disease enhanced tolerance, consistent with bilirubin’s direct antiplasmodial activity [23].

Dysregulation of glucose metabolism in *Pcc*-infected *Blvra^G17A^*, *Blvra^E97A^* and *Blvra^-/-^* mice (Fig. 6I-L) suggests an additional dimension distinct from weight loss and thermoregulation trajectories. While *Blvra^WT^* mice maintained euglycemia (∼100-150 mg/dL) throughout the entire course of infection, *Blvra* mutants developed severe hyperglycemia (∼400-600 mg/dL), with *Blvra^E97A^* mice showing the most pronounced response despite retaining BVRA protein expression (Fig. 6I-L). This suggests that sub-threshold accumulation of unconjugated bilirubin not only fails to control parasitemia but may exacerbate metabolic dysregulation. The hyperglycemic phenotype likely reflects multiple converging mechanisms centered not only on inhibition of parasite consumption via glycolysis [23] but possibly on infection-driven metabolic stress including dysregulated hepatic glucose output [30] and impaired peripheral glucose utilization driven by inflammatory mediators released during uncontrolled parasitemia. This finding has implications for therapeutic strategies, as interventions must address not only parasite clearance through bilirubin-mediated resistance but also the metabolic consequences of failed resistance, including thermoregulatory collapse and glucose dysregulation. The threshold phenomenon thus extends beyond parasite control to encompass maintenance of metabolic homeostasis, reinforcing that 20-30 μM bilirubin represents the minimum concentration required for integrated host defense encompassing both antiparasitic activity and preservation of physiological stability. For therapeutic strategies, these findings indicate that such interventions must achieve sustained plasma unconjugated bilirubin concentrations exceeding 20-30 μM, as measured by UnaG or equivalent methods that account for albumin binding during the acute parasite clearance phase. This has implications for patient stratification and therapeutic monitoring, as accurate bilirubin quantification using UnaG-based or equivalent assays (rather than conventional clinical methods) will be essential for determining whether protective thresholds are achieved .

Several considerations qualify these conclusions. The protective threshold was established in the *Pcc* AS model, and whether identical thresholds apply to other *Plasmodium* species or more virulent strains requires confirmation. The mechanisms linking bilirubin’s multiple effects on mitochondrial function, hemozoin formation, and food vacuole integrity remain incompletely defined, particularly whether these are independent or functionally coupled processes. Future studies should employ UnaG-based quantification systematically across multiple *Plasmodium* species and strains, expand human cohorts with comprehensive bilirubin measurements stratified by disease severity, and investigate whether threshold concentrations vary by host factors. In conclusion, converging evidence from genetic, biochemical, and pharmacological approaches, unified by UnaG-based quantification methodology, establishes that unconjugated bilirubin concentrations of 20-30 μM represent the minimum threshold for antimalarial protection, providing a quantitative framework for therapeutic development and biomarker-guided treatment strategies.

## Materials and Methods

### Mice

Mice were bred and maintained under specific pathogen-free (SPF) conditions at the Gulbenkian Institute for Molecular Medicine (GIMM). Mice were housed at standard *vivarium* temperature (22°C) in a 12-hour light/dark cycle with free access to water and standard chow pellets. All experimental protocols were approved in a two-step procedure, by the Animal Welfare Body of the IGC and by the Portuguese National Entity that regulates the use of laboratory animals in research (Direção Geral de Alimentação e Veterinária; DGAV). Experimental procedures followed the Portuguese (Decreto-Lei no 113/2013) and European (Directive 2010/63/EU) legislation. C57BL/6J mice were obtained from the GIMM animal facility. C57BL/6J *Blvra*^−/−^ mice were generated at Ozgene (Australia), as described [6]. C57BL/6J *Blvra^G1^*^7A^ and C57BL/6J *Blvra*^E97A^ mouse strains were developed in collaboration with the GIMM’s transgenic unit. Briefly, they were generated by CRISPR/Cas9, using gRNAs targeting the GTGGTTGGTGTTGGCAGAGC or the ATGCCATGGGGTATTCCACG sequence respectively for the C57BL/6J *Blvra^G17A^* and the C57BL/6J *Blvra^E97A^* mice, and the replacement oligonucleotides CTCCATTGCATAGCTGGGCTGTTTTTAAACCCCACATCGGTCTTTGATATTTCAGCCAAAGAGGAAATTTGGTGTGGTAGTGGTTGGTGTGGCACGGGCTGGCTCTGTGAGGATAAGGGACTTGAAGGATCCACACTCTTCAGCATTCCTAAACCTGATTGGATATGTGTC or TTTGAAGTGTCCTTTCCCTGTCTTTTTGTTTTCAAAGGCAGTTTCTTCAGGCTGGCAAGCATGTCCTCGTCGCCTATCCCATGGCATTGTCATTTGCGGCAGCGCAGGAGCTGTGGGAGCTGGCTGCACAGAAAGGTGATGTT respectively for the C57BL/6J *Blvra^G17A^* and C57BL/6J *Blvra^E97A^*mice (all oligos purchased from Integrated DNA Technologies). This resulted in the replacement of glycine 17 for an alanine, in C57BL/6J *Blvra^G17A^*, and glutamate 97 for an alanine, in C57BL/6J *Blvra^E97A^*.

For better discrimination of *Blvra^WT^ vs. Blvra^G17A^* strains during genotyping we replaced in the *Blvra^G17A^* strain, the 5’ GTT and 3’AGA codons, flanking the mutated GGC->GCA codon, by the synonymous 5’ GTG and 3’CGG codons, respectively. For better discrimination of *Blvra^WT^ vs. Blvra^E97A^* strains during genotyping we replaced in the *Blvra^E97A^* strain, the 5’GTG and 3’TAC codons, flanking the mutated GAA->GCC codon, by the synonymous 5’GTC and 3’TAT codons, respectively.

Ribonucleoprotein complexes (1μM gRNA, 100 ng/μL Cas9 protein) were mixed with the replacement oligo (30 ng/μL) and microinjected into the pronucleus of fertilized C57BL/6J mouse oocytes, which were transferred into the uterus of pseudo pregnant females, according to standard procedures. Genotyping was performed by PCR on genomic DNA using primers to amplify WT or mutant sequences. Sanger sequencing revealed 2 C57BL/6J *Blvra^G1^*^7A^ founders and C57BL/6J *Blvra^E97A^*3 founders. These were bred with C57BL/6J mice to obtain heterozygous mice for the desired mutations. Offspring was genotyped using the folliwng primer pairs: *Blvra^WT^*: *Fwd* TAGTGGTTGGTGTTGGCAGA; *Rev*: GGCTTTCCCTTACTCTGGGTC; *Blvra^G17A^*: TAGTGGTTGGTGTTGGCAGA; *Rev*: GGCTTTCCCTTACTCTGGGTC; *Blvra^E97A^ Fwd*: AGCATGTCCTCGTGGAATAC, *Rev*: TGGGAACTCAGTAGCAAAGCC. The progeny of positive mutant lines and C57BL/6J mice was subsequently bred to homozygosity.

### RNA extraction and RT-qPCR

Mice were sacrificed by CO_2_ inhalation at steady state, transcardially perfused *in toto* with ice-cold PBS (1X, 20 mL) and organs were harvested, snap frozen in liquid nitrogen and stored at -80°C. Total RNA was extracted using tripleXtractor reagent (GRISP), chlorophorm, isopropanol and ethanol, according to manufacturer’s instructions. cDNA was synthesized using the Xpert cDNA Synthesis Mastermix (GRiSP), followed by RT-qPCR using the iTaq Universal SYBR Green Supermix (Bio-Rad) on a QuantStudio™ 7 Flex Real-Time PCR System (Applied Biosystems). Transcript values were calculated from the threshold cycle (Ct) of each gene using the 2^-ΔCT^ method using Acidic ribosomal phosphoprotein P0 (*Rplp0*) as the housekeeping control gene. Primers for qPCR include: *Rplp0*, Fwd: 5’-CTTTGGGCATCACCACGAA-3’, Rev: 5’-GCTGGCTCCCACCTTGTCT-3’; *Blvra*, Fwd: 5′-AGCCGCTGGTAAGCTCC-3′, Rev: 5′-ACCAACCACTACCACACCAAA-3′.

### BVRA Enzymatic Activity Assay

BVRA enzymatic activity was measured as described [37]. Snap-frozen tissues were homogenized in homogenization buffer (10 mM Tris-HCl pH7.5, 250 mM sucrose) and the enzymatic activity was then measured in 50 µL of protein homogenate in assay buffer (100 mM Tris-base, 1 mM EDTA, pH 8.7, 1 mM NADPH, and 3 µM biliverdin), at 37°C in a spectrophotometer with shaking for 1hr. The rate of reaction was determined by monitoring the change in absorbance at 450 nm over time. Bilirubin levels in the assay samples were determined using a standard curve of bilirubin IXα (0-300 µM) (Frontier Chemicals) and then normalized to cellular protein levels and presented as bilirubin concentration per ug of total protein. The activity assay was performed in a final volume of 600 µL.

### Western Blot

Mice were sacrificed, transcardially perfused *in toto* with ice-cold PBS (1X, 20 mL) and organs were harvested and kept at -80°C until tissue lysates preparation. Tissues were lysed using 2% SDS-PAGE sample buffer (100mM Tris, pH 6.8, 20% glycerol, 4% SDS, 0.2% bromophenol blue, 100mM DTT and 1X protease inhibitor cocktail (cOmplete™, Mini, EDTA-free Protease Inhibitor Cocktail; Roche)) or NP40 extraction buffer (0,15M NaCl, 1% NP-40, 0,05M Tris, 1X protease inhibitor cocktail (cOmplete™, Mini, EDTA-free Protease Inhibitor Cocktail; Roche) and homogenized in a tissue lyser (Qiagen) with tungsten carbide beads (Qiagen). For the SDS-PAGE sample method, supernatants were collected and total protein was quantified at λ_280_nm using the DS-11 FX Spectrophotometer (DeNovix). For NP40 method, supernatant was collected and protein was quantified via Bradford assay (BioRad, Cat#5000006). Protein was resolved (50µg) on a 12% SDS-PAGE and transferred to Polyvinylidene fluoride (PVDF) membranes. Membranes were blocked for 1 hour at room temperature (5% bovine serum albumin in 1X TBS-T), washed in 1X TBS-T and incubated with primary antibodies, overnight at 4°C. The primary antibodies used were rabbit polyclonal anti-BVRA (Thermofisher scientific, PA5-92059; 1:750) and rabbit polyclonal anti-β-actin (Cell Signaling Technology, 4967; 1:1000). Membranes were washed 3 times (1X TBS-T) and incubated (1 h; RT) with the peroxidase-conjugated secondary antibody (HRP conjugated goat anti-rabbit IgGH+L; Invitrogen, #31460; 1:5000). Membranes were washed 3 times (1X TBS-T) and peroxidase activity was detected using SuperSignal™ West Pico PLUS Chemiluminescent Substrate (ThermoFisher Scientific). Blots were developed using Amersham Imager 680 (GE Healthcare), equipped with a Peltier cooled Fujifilm Super CCD. Densitometry analysis was performed using ImageLab software, from images without saturated pixels.

### *Plasmodium chabaudi chabaudi* AS infection and disease assessment

Mice (females and males, 10-16 weeks old) were infected with *Plasmodium chabaudi chabaudi* AS transgenic GFP-expressing *Pcc* AS (Pcc AS-GFPML)[25]. Infections were performed by intraperitoneal (i.p.) administration of freshly isolated blood (Passage 29; 2 × 10^6^ infected RBC diluted in 200 μL PBS) collected from a previously infected C57BL/6J mouse. Mice were monitored daily from day 0 (day of infection) onwards for parasitemia (% infected RBC; iRBC), parasite burden (number of iRBC per μL of blood), body weight (Ohaus CS200 scaler, Sigma Aldrich), core (i.e. rectal) body temperature (Rodent thermometer; BIO-TK8851, Bioset), blood glucose concentration (AccuCHECK Performa glucometer, Roche) and survival, essentially as described [38]. Briefly, the number of RBC per μL of blood was quantified by flow cytometry (LSR Fortessa X20 analyzer; BD Bioscience) using a standard concentration of reference latex beads (10 μm; Coulter CC Size Standard L10, Beckman Coulter, no. 6602796), gating on RBC, based on size and granularity and on bead population.

### UnaG protein synthesis and purification

UnaG protein synthesis and purification was done as before [6]. Briefly, UnaG was expressed in pMAL-6P2-6xHIS in BL21(DE3) cells [6]. Starter cultures were grown to saturation in Luria Broth (LB) (overnight at 37°C), diluted 10-fold in LB and grown to an OD600 of 0.3 at 37°C, after which cultures were moved to 18°C. UnaG expression was induced at OD600 of 0.6, by the addition of 400 μM isopropyl β-D-1thiogalactopyranoside (16 hours, 18°C). Cells were harvested by centrifugation (3,800 g, 4°C, 25 min), resuspended in 15 ml of resuspension buffer (50 mM HEPES, 300 mM NaCl, 0.5 mM TCEP, 10% glycerol, 1 mM PMSF, 2.34 μM leupeptin, 1.45 μM pepstatin at pH 7.4) per liter of culture. The sample was further lysed by sonication (QSonica Sonicator) with an amplitude of 5% in a pulse mode of 0.8sec ON and 0.5sec OFF for total 30 s. Lysate was clarified by centrifugation (26,000 g, 4°C, 30 min) and loaded onto an amylose column (MBPTrap HP Column, GE Healthcare). Protein was eluted with 20 mM maltose in protease buffer (50 mM Tris, 150 mM NaCl, 0.5 mM TCEP, 10% glycerol, 0.01% TritonX-100 at pH 7.4). MBP was removed by the addition (16 hours, 4°C) of PreScission Protease (GE Healthcare). UnaG was purified via a nickel column (HisTrap HP Column, GE Healthcare) to remove the cleaved tag and eluted with (30 mM Tris, 1M NaCl, 0.5 mM TCEP, 10% glycerol, 500 mM imidazole, pH 7.4). UnaG was further purified on a gel-filtration column (S-200, GE Healthcare) in gel filtration buffer (50 mM HEPES, 300 mM NaCl, 0.5 mM TCEP, 10% glycerol at pH 7.4), concentrated using 10 kDa amicon ultracentrifugal filter (Merck), and flash frozen in gel filtration buffer at 30% glycerol for storage at −80°C.

### UnaG-based assay for quantification of unconjugated bilirubin

Blood from *Blvra*^+/+^, *Blvra*^-/-^, *Blvra^G17A^* and *Blvra^E97A^* mice was obtained from the tail vein, before (Day 0) and during *Pcc* infection. Samples were collected and immediately centrifuged (1000 g, 15 min) and plasma was collected, frozen in liquid nitrogen and stored at −80°C until used for bilirubin quantification. Total plasma unconjugated bilirubin was determined using UnaG [39]. Plasma samples from mice noninfected or infected with *Pcc* was diluted 1:100 in PBS. Then, 100 pM of UnaG was added to the diluted plasma sample and incubated for 10 min at RT. After incubation, fluorescence intensity was measured at excitation λ480 nm and emission λ530 nm using a microplate reader (Promega GloMax). Bilirubin IXα (Frontier Chemicals) was used as standard.

### Statistical analysis

Statistically significant differences between more than two groups were assessed using two-way ANOVA with Tukey’s multiple comparison test or two-way ANOVA with Linear Mixed Effects Model. Survival curves are represented by Kaplan-Meier plots and differences between the groups were assessed using the log-rank test. All statistical analyses were performed using GraphPad Prism 10 software. Differences were considered statistically significant at a *P* value <0.05. NS: not-significant, *p*>0.05; \**p*<0.05; \*\**p*<0.01; \*\*\**p*<0.001; \*\*\*\**p*<0.0001.

## Declaration of generative AI and AI-assisted technologies in the manuscript preparation process

During the preparation of this work the author(s) used [Claude] in order to [edit the manuscript body of text]. After using this tool/service, the author(s) reviewed and edited the content as needed and take(s) full responsibility for the content of the published article.

## Acknowledgments

The authors are indebted to all members of the Inflammation group (GIMM) for insightful technical and intellectual contributions, to the staff at the GIMM flow cytometry and animal facility.

## Funding

Fundação para a Ciência e Tecnologia (UI/BD/152257/2021 to MM, 2020.04797.BD and COVID/BD/153665/2024 to AF; FEDER/29411/2017 to SR; PTDC/MED-FSL/4681/2020 DOI 10.54499/PTDC/MED-FSL/4681/2020 to SP; CEECIND/01589/2017 and DOI 10.54499/2023.11177.PEX to ARC; 2023.09168.CEECIND to EJ; FEDER/29411/2017, PTDC/MED-FSL/4681/2020 DOI 10.54499/PTDC/MED-FSL/4681/2020, 2022.02426.PTDC DOI 10.54499/2022.02426.PTDC and Congento LISBOA-01-0145-FEDER-022170 to MPS.

Gulbenkian Foundation (SR, SC, MPS and IBB 2021-51/BI-D/2021 to ST).

GIMM Foundation (GIMM/BI/36-2025 to MM, GIMM/BI/37-2025 to ST, GIMM Cross-Site collaborative project to MPS).

la Caixa Foundation HR18-00502 (EJ, MPS).

American Heart Association/Paul Allen Frontiers Group (Project 19PABH134580006 to BDP).

DFG Cluster of Excellence “Balance of the Microverse” EXC 2051; 390713860 (E.J., M.P.S. as associated member)

NIH/NIA (1R21AG073684-01, R01AG071512 to BDP).

The Johns Hopkins Catalyst Award (BDP).

Solve ME/CFS Initiative (Grant 90089823 to BDP). US Public Health Service (Grant DA044123 to BDP).

Oeiras-ERC Frontier Research Incentive Awards (MPS).

H2020-WIDESPREAD-2020-5-952537 SymbNET Research Grants (MPS)

## Author Contributions

Conceptualization: MPS and MM

Formal analysis: MM and AF

Resources: BDP

Investigation: AF, AN, ARC, MM, MM, STR, SR, SP, SC

Visualization: MM and MPS

Funding acquisition: MPS

Project administration: MPS

Supervision: MPS and EJ

Writing – original draft: MPS

Writing – review & editing: MM

